# Effects of time and isolation on plant diversity: testing island biogeography theory with an eco-evolutionary model

**DOI:** 10.1101/100289

**Authors:** Juliano Sarmento Cabral, Robert J. Whittaker, Kerstin Wiegand, Holger Kreft

## Abstract

**Aims:** The General Dynamic Model of oceanic island biogeography (GDM) predicts how biogeographical rates, species richness, and endemism vary depending on island age, area, and isolation, based on the interplay of colonization, extinction, and speciation. Here, we used a simulation model to test whether GDM predictions may arise from individual- and population-level processes.

**Location:** Hypothetical hotspot islands.

**Methods:** Our model (i) considers an idealized island ontogeny, (ii) metabolic constraints, and (iii) stochastic, spatially-explicit, and niche-based processes at the level of individuals and populations (plant demography, dispersal, competition, mutation, and speciation). Isolation scenarios involved varying dispersal ability and distances to mainland.

**Results:** Humped temporal trends were obtained for species richness, endemic richness, proportion of cladogenetic endemic species, number of radiating lineages, number of species per radiating lineage, and biogeographical rates. The proportion of anagenetic endemics and of all endemics steadily increased over time. Extinction rates of endemic species peaked later than for non-endemic species. Species richness and the number of anagenetic endemics decreased with isolation as did rates of colonization, anagenesis, and extinction. The proportion of all endemics and of cladogenetic endemics, the number of cladogenetic endemics, of radiating lineages, and of species per radiating lineage, and the cladogenesis rate all increased with isolation.

**Main conclusions:** The results confirm most GDM predictions related to island ontogeny and isolation, but predict an increasing proportion of endemics throughout the experiment: a difference attributable to diverging assumptions on late island ontogeny. New insights regarding the extinction trends of endemics further demonstrate how simulation models focusing on low ecological levels provide tools to test biogeographical-scale predictions and to develop more detailed predictions for further empirical tests.

## INTRODUCTION

Geographical isolation is considered one of the key drivers of species diversification in both insular (e.g. Heaney, 2000; Whittaker & Fernández-Palacios, 2007; Rosindell & Phillimore, 2011) and continental systems (e.g. Linder, 2005; Rieseberg & Willis, 2007; Pennington *et al.*, 2010). Mechanisms by which isolation operates to promote divergence and hence diversification include founder effects and genetic drift via limited (or no) gene exchange with other populations (non-adaptive speciation; Rundell & Price, 2009). Isolated populations may be further subject to differential selective pressures compared to the source areas, triggering adaptive (e.g. ecological) speciation (Rundell & Price, 2009). These mechanisms are characteristic of oceanic islands, causing high numbers of endemic species (Whittaker & Fernández-Palacios, 2007; Steinbauer *et al.*, 2012) and providing model systems for investigating isolation effects (MacArthur & Wilson, 1967; Weigelt & Kreft, 2013; Warren *et al.*, 2015). Island biogeography theory holds that isolation reduces colonization rates and the species richness at the dynamic equilibrium between colonization, speciation, and extinction, while increasing the relative contribution of speciation to species richness (MacArthur & Wilson, 1967; Whittaker *et al.*, 2008). These effects on speciation have been formalized within the General Dynamic Model of oceanic island biogeography (GDM), the distinguishing feature of which is to posit that diversity patterns within and across archipelagos are influenced in a predictable fashion by the geodynamics of oceanic islands over their lifespan (i.e. their ontogeny, Whittaker *et al.*, 2008, 2010).

For classic hotspot islands, the GDM predicts humped trends in richness and endemism as a function of the rise and decline of island area, elevation, and habitat heterogeneity over the island’s life span (Whittaker *et al.*, 2008, 2010). Throughout these environmental changes along the island’s life span, isolation affects biogeographical patterns by changing the amplitude but not the shape of the temporal trends (Whittaker *et al.*, 2008). More recently, Borregaard *et al.* (2016b) used a simulation model to explore GDM properties, in particular examining isolation effects, and alternative island ontogenies. These simulations supported the internal logic of the GDM, and provided additional island age-related predictions. For instance, more isolated islands had lower colonization and extinction rates, as well as higher speciation rates (Borregaard *et al.*, 2016b). Consequently, isolation should decrease overall species numbers, but increase the number and proportion of endemic species. The main explanation is that on remote islands radiating lineages fill niches that would be occupied by colonist species on less isolated islands (Heaney, 2000; Whittaker *et al.*, 2008). In sum, the capacity to generate explicit biogeographical predictions has made the GDM an important framework for many recent island studies (Borregaard *et al.*, 2016a).

A general limitation to testing GDM predictions is that systematic empirical data for islands over their entire ontogeny cannot be obtained due to the protracted nature of volcanism and erosion processes, which destroy fossils. The intractability of measuring colonization, extinction, and diversification histories has justified the use of space-for-time substitutions involving analyses of islands of different ages within archipelagos as surrogates for following the long-term dynamics of single islands (Borregaard *et al.*, 2016a). Alternatively, the application of mechanistic simulation models to investigate GDM predictions has been identified as a particularly promising avenue (Borregaard *et al.*, 2016a). Such models have full control of relevant factors and processes and thus exclude confounding effects. Among these are several spatial processes and factors not fully described in the theory, such as island hopping followed by parallel radiations (Whittaker & Fernández-Palacios, 2007; Losos & Ricklefs, 2010), rescue effects (Brown & Kodric-Brown, 1977), intra-archipelagic spatial settings (Cabral *et al.*, 2014, Weigelt *et al.*, 2016), and the eco-evolutionary history specific to each island, archipelago, and taxon (Whittaker & Fernández-Palacios, 2007; Bunnefeld & Phillimore, 2012). The variety of mechanisms contributing to isolation – from distance to potential source pools to varying dispersal ability of different taxa–also opens up alternative means of quantifying (Weigelt & Kreft, 2013) and simulating isolation in mechanistic models. For example, assemblage-level models can simulate single islands that vary only in distance to the mainland but that have identical environmental settings and geological trajectory (e.g. Borregaard *et al.*, 2016b). In previous work, Cabral *et al.* (submitted), present a BioGeographical Eco-Evolutionary Model (BioGEEM) consistent with and supporting GDM temporal predictions of species richness and biogeographical rates via population-level processes. However, only one isolation scenario was investigated by Cabral *et al.* (submitted) and thus GDM predictions of isolation effects remain to be investigated in more details by such trait-based, spatially-, and demographically-explicit models. This is the purpose of the present study.

BioGEEM is niche-based and integrates metabolic, demographic, dispersal, and competition constraints at the local scale (and at the level of individuals and populations), with evolutionary and environmental processes at biogeographical scales (Cabral *et al.*, submitted). The model is specified for terrestrial seed plants and has a hierarchical structure that links ecological and evolutionary processes to local temperature and body mass via metabolic trade-offs based on the metabolic theory of ecology (Brown *et al.*, 2004). As a result, all biogeographical patterns emerge from processes operating at local scales and low levels of ecological organization, e.g. individual dispersal, resource competition, and local population dynamics. The model properties of BioGEEM thus differs from previous island models, which concentrate on geologically static islands (Kadmon & Allouche, 2007; Hortal *et al.*, 2009; Rosindell & Phillimore, 2011; Rosindell & Harmon, 2013), do not incorporate evolutionary processes (Kadmon & Allouche, 2007; Hortal *et al.*, 2009; Rosindell & Harmon, 2013), simulate ecologically neutral processes (Rosindell & Phillimore, 2011; Rosindell & Harmon, 2013; Valente *et al.*, 2014, 2015; Borregaard *et al.*, 2016b), and/or are spatially-implicit (Valente *et al.*, 2014, 2015; Borregaard *et al,* 2016b). In short, BioGEEM includes a combination of properties and assumptions not yet applied for assessing the eco-evolutionary logic of GDM predictions, which might be useful for the derivation of generalizable insights (Evans *et al.*, 2013; Cabral *et al.*, 2016; see Cabral *et al.*, submitted). To achieve this, we integrated into BioGEEM the GDM assumptions of geologically dynamic islands, habitat heterogeneity, and niche-based adaptive radiations. In our simulation experiment, we tested GDM-based hypotheses related to eight biogeographical variables (Table 1) by varying island isolation in two ways: via dispersal ability of the species source pool and via distance to the mainland.

**Table 1.** Hypotheses based on the GDM (Whittaker *et al.*, 2008; Borregaard *et al.*, 2016b) for eight biogeographical variables and the model output used for their evaluation. The BioGEEM generates time-series of all variables covering the entire lifespan of islands. We adopted the simplest method for calculating the rates to make these comparable to GDM predictions, given as the number of events occurring within arbitrary time intervals.

| Variable | Model output | Hypotheses |  |
| --- | --- | --- | --- |
|  |  | Temporal trend | Isolation effects |
| 1. Species richness | Number of species | humped | negative |
| 2. Endemic richness | Number of anagenetic and cladogenetic species | humped | positive |
| 3. Proportion of endemics | Percentage of all endemic, only anagenetic, only cladogenetic species richness in relation to all species | humped | positive |
| 4. Radiating lineages | Number of lineages showing cladogenesis | humped | positive |
| 5. Radiation extent | Number of species per radiating lineage | humped | positive |
| 6. Colonization rate | Number of colonization events per time interval | humped | negative |
| 7. Extinction rate | A. Number of all extinction events per time interval | humped | negative |
|  | B. Number of extinction events of endemic species per time interval | humped | negative |
| 8. Speciation rate | Number of anagenetic and cladogenetic speciation events per time interval | humped | positive |

## MATERIALS AND METHODS

### Modelling approach

We briefly summarize the model below. A detailed description and parameter settings can be found in Cabral *et al.* (submitted) and in Appendix S1 in Supporting Information.

### State variables and scales

The model is grid-based (Fig. 1a), with a cell size of 1 km^2^. Each island cell was assigned to an elevational level and mean annual temperature (the lowest elevation band was assigned 25 °C). The model agents are stage-structured plant populations (seeds, juveniles, and adults), given in number of individuals. Populations belong to species, defined by combinations of autecological attributes (hereafter: species properties): environmental requirements (maximum cell suitability, optimum temperature, temperature tolerance, optimum island side, and island side tolerance), short- and long-distance dispersal abilities, Allee threshold, body sizes (seed, juvenile and adult body sizes), and phenological ordering. Habitat requirements depict preferences associated with elevation (i.e. temperature) as well as wind and precipitation patterns in islands (i.e. island side). Body mass and local temperature determine all demographic transitions, mutation rates, the space exploited by an individual, carrying capacity, and time for speciation. These metabolic constraints account for increasing metabolism with temperature and decreasing metabolic rate with body mass (Brown *et al.*, 2004). Demographic transitions are germination, sexual maturation, reproduction, and density-independent mortality. A cell can hold one population per species, but as many species as there is space available. Consequently, species assemblages emerge from local resource competition among populations (Cabral & Kreft 2012; Cabral *et al.*, submitted). The state variables comprise the spatial distribution of seed, juvenile, and adult abundances of each species and the unoccupied area. Each time step represents one year and a complete simulation runs over 2.21 million time steps.

**Figure 1.**
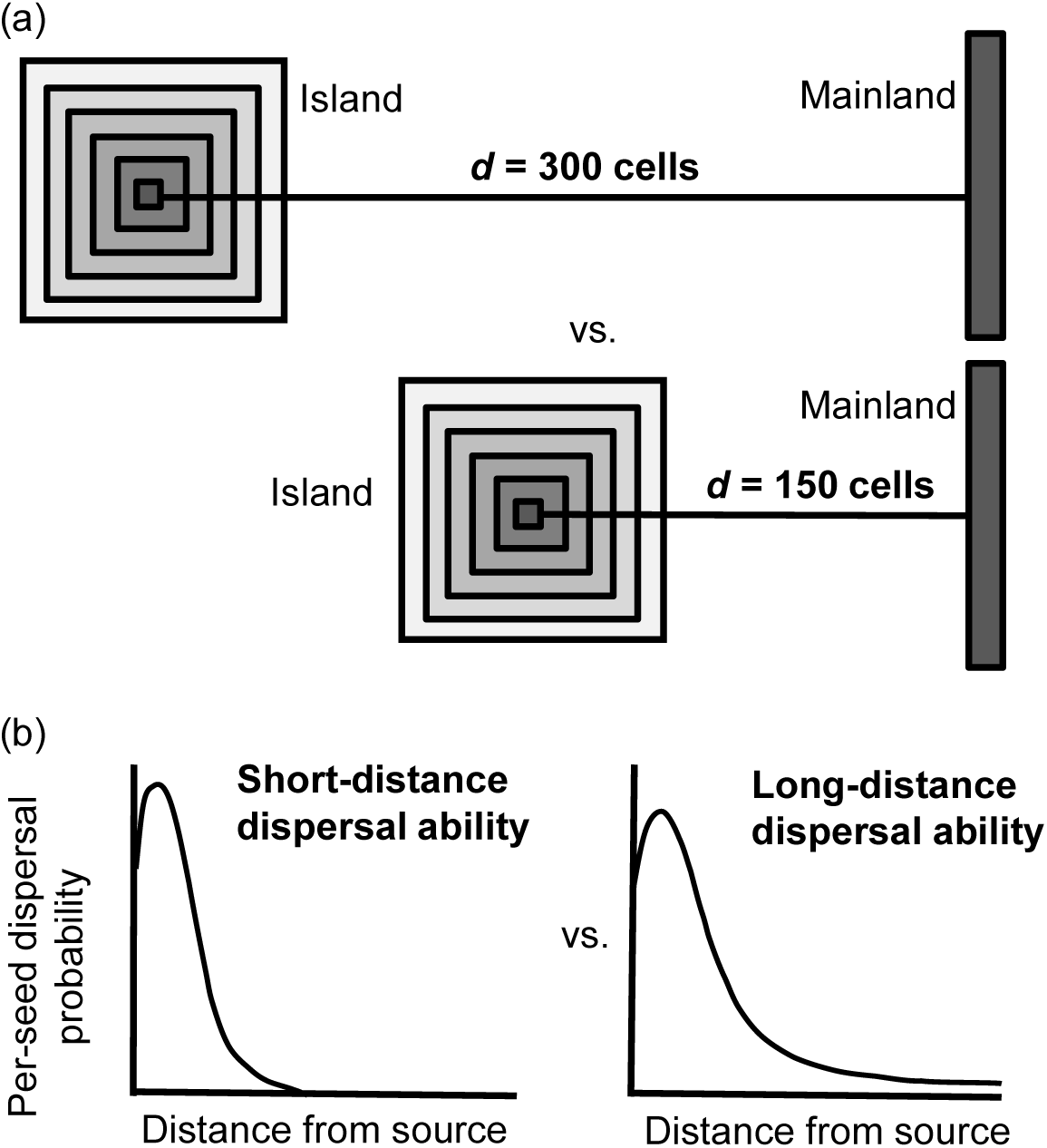
Isolation scenarios. (a) Distance scenarios based on the distance *d* from the island centre to the mainland: *d*=300 vs. *d*=150 cells. (b) Distance scenarios based on the dispersal ability of the mainland source pool of species: short-distance vs. long-distance dispersal ability (thin and fat kernel tails, respectively).

### Initialization

Simulations are initialized with a species pool size of 1000 species, and the intervals for randomly drawing the species properties (see Appendix S1). Species properties vary from representing specialists (e.g. narrow temperature amplitudes) to generalists (e.g. all temperatures and island sides). For each species, the habitat suitability matrix, ***H***, is initialized based on the species′ environmental requirements. A species-specific dispersal kernel ***D*** is initialized as a twodimensional, grid-based Clark’s 2Dt kernel, with two parameters, α and p, which describe short- and long-distance dispersal, respectively (Fig. 1b; Clark *et al.*, 1999; Nathan & Muller-Landau, 2000). The stage-specific abundance matrices (seeds, juveniles, and adults) and the matrix with the area occupied by all individuals are initialized empty.

### Submodels

At each time step, a series of submodels are executed in the following order: dispersal from mainland, population update 1, reproduction, intra-island dispersal, mutation, speciation, population update 2, and environmental dynamics. In each submodel, the state variables of each species are updated following the species phenological ordering:

#### Dispersal from mainland

A random number of seeds per mainland cell from ten random mainland species are dispersed to the island according to ***D***.

#### Population update 1

Abundance matrices are updated by: A) turning juveniles to adults, B) applying density-independent mortality to remaining juveniles, C) germinating seeds, and D) applying seed mortality.

#### Reproduction

If adults are present, the number of seeds produced by each species in each cell is given by the Beverton-Holt reproduction function, extended with Allee effects (Cabral & Schurr, 2010).

#### Intra-island dispersal

The produced seeds are dispersed within the island following ***D***.

#### Mutation

As a previous step to cladogenesis (see next submodel), each seed dispersed is randomly assigned as mutant given a metabolic-constrained probability (Brown *et al.*, 2004) via point-mutation (Rosindell & Phillimore, 2011). Mutant seeds received random properties according to phylogenetic constraints (values within ±50% of ancestral values). The ***H***, ***D***, and abundance matrices for these mutant individuals are initialized (see ‘*Initialization*’).

#### Speciation

Two modes of speciation are considered: anagenesis (differentiation from mainland species) and cladogenesis (within-island diversification). The submodel checks whether enough time has passed to update mutant individuals (cladogenesis) or colonizers (anagenesis) as a distinct species (i.e. 'protracted speciation' _ Rosindell & Phillimore, 2011). The time for speciation is species-specific and follows metabolic constraints to account for longer generations of larger species (Brown *et al.*, 2004). Anagenesis could be delayed due to gene flow from the mainland.

#### Population update 2

After species status update, the submodel applies density-independent mortality to adults and updates the seed bank.

#### Environmental dynamics

Environmental events occur that mimicked the geological trajectory of a typical oceanic island (Whittaker & Fernández-Palacios, 2007; Whittaker *et al.*, 2008), namely island growth due to volcanic activity followed by a slower erosion-dominated phase. The island grows and shrinks by gaining or losing belts of cells, respectively, and by increasing or decreasing elevation and thus local temperature accordingly. Islands grow every 0.13 Ma, whereas erosion takes place every 0.26 Ma (see Appendix S1). After every environmental change event, ***H*** is recalculated for every species.

### Output

The model records time-series of species richness (total, anagenetic, and cladogenetic endemics), number of endemic lineages (including species that evolved from the same ancestor), and number of species per endemic lineage, as well as the number of colonization, speciation, and extinction events.

### Study design

All intervals for drawing species properties, the model, and the scenario specifications are provided in Appendix S1. To test our hypotheses (Table 1), we set up four isolation scenarios (Fig. 1). The scenarios encompassed a full-factorial design, varying the shortest distance between the island at maximum size and the mainland (150 vs. 300 cells), as well as the dispersal ability of the mainland species pool (high vs. low long-distance dispersal ability). Greater long-distance dispersal ability (*p*_high_) was obtained by systematically varying the dispersal parameter *p* for all species of the mainland source pool from the scenario with low long-distance dispersal ability (*p*_low_): *p*_high_ = *p*_low_– 0.2 (*p*_low_ values in Appendix S1). Note that although 150-or 300-cells distance might seem close to the mainland, isolation was assured by generally low long-distance dispersal ability. Nevertheless, we calibrated the dispersal ability so that even the most isolated islands could receive at least one species during time steps with minimum island size. All islands had a maximum size of 11 × 11 cells.

The simulation experiment comprised 20 replicate runs per isolation scenario, with each replicate having a different species pool to represent different mainland species compositions. Outputs for each time step were averaged over replicates (only averaged time-series are shown for visual clarity, but see Figure S1 of the Appendix S2 for examples of 95% standard deviation envelopes around average time-series). To make results comparable to GDM predictions, colonization, speciation, and extinction rates were calculated by summing the number of colonization, speciation, or extinction events within time intervals of 0.01 Myr. Only successful colonization, i.e. germination and establishment, was considered.

## RESULTS

Total species richness and endemic richness showed a clear humped relationship with island age (Fig. 2). Richness peaks lagged behind maximum island size – a pattern that was most pronounced for more isolated islands (Fig. 2) and for cladogenetic endemics (Fig. 2b). Except at the very final island stages where species numbers converged, more isolated islands had the lower total species and anagenetic endemic richness and the higher cladogenetic endemic richness (Fig. 2).

**Figure 2.**
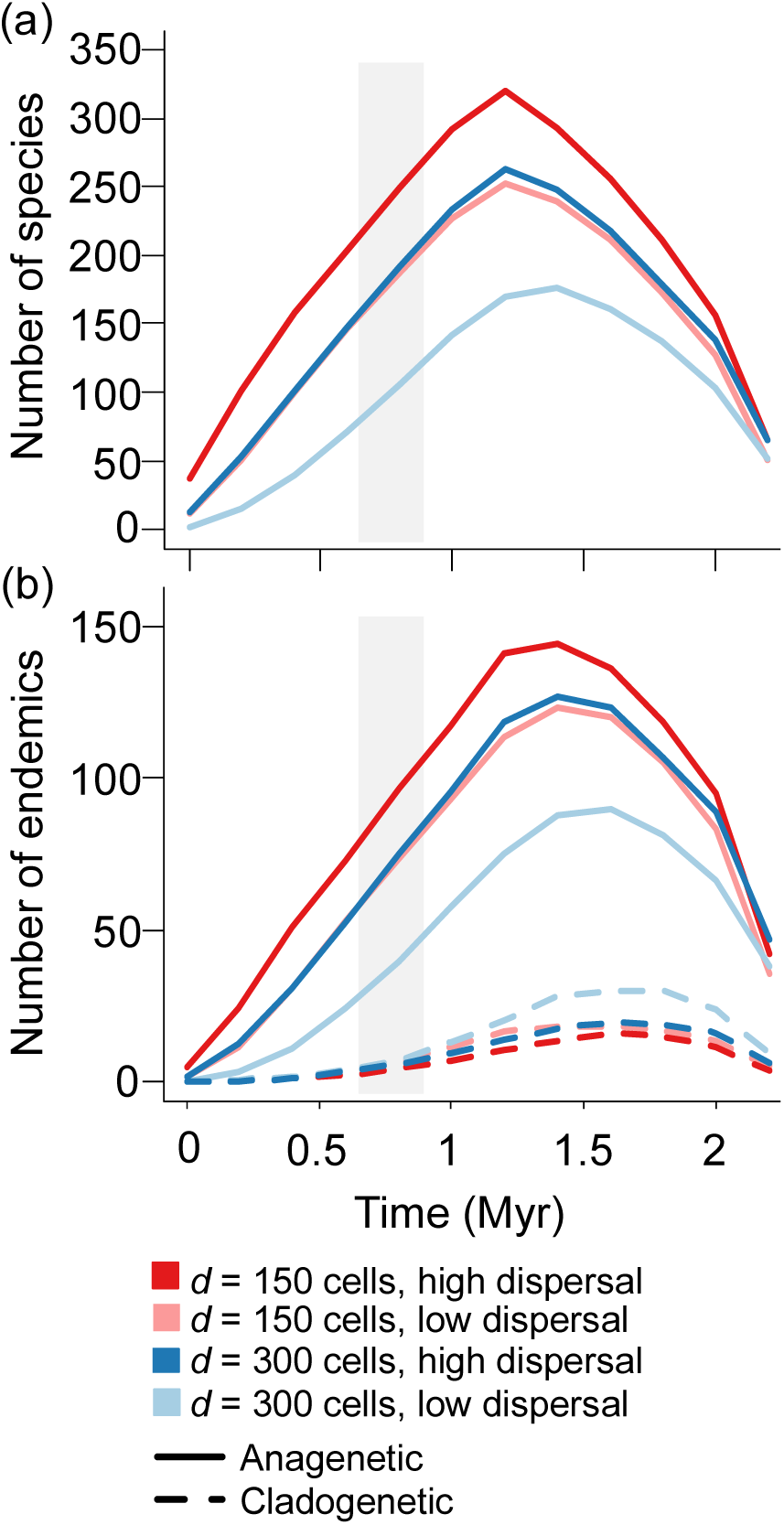
Temporal trends in species numbers of four different isolation scenarios. Time-series of: (a) number of all species; and (b) number of anagenetic and cladogenetic endemics. Isolation scenarios were given by changing the distance *d* from mainland and long-distance dispersal ability of the source pool. Time-series were averaged within environmental time steps and over 20 replicate runs. The shaded area indicates the period with maximum island size. Note in (a) that two intermediate scenarios (*d* = 150 cells, low dispersal and *d* = 300 cells, high dispersal) are barely distinguishable.?

The proportion of endemics increased over time and with isolation (Fig. 3a). This increase was mostly driven by anagenetic endemics and there was little difference among isolation scenarios (Fig. 3b). In contrast, the proportion of cladogenetic endemics showed a shallow, humped relationship with island age and more isolated islands attained higher proportions (Fig. 3b). The number of radiating lineages also exhibited a humped relationship with island age. More isolated islands had higher values, and there was a strong difference between isolation scenarios at advanced island age (Fig. 3c). The number of species per radiating lineage showed a general humped relationship with island age, and varied less clearly with isolation, with only the islands isolated by both distance and dispersal having evidently higher values than the other islands (Fig. 3d).

**Figure 3.**
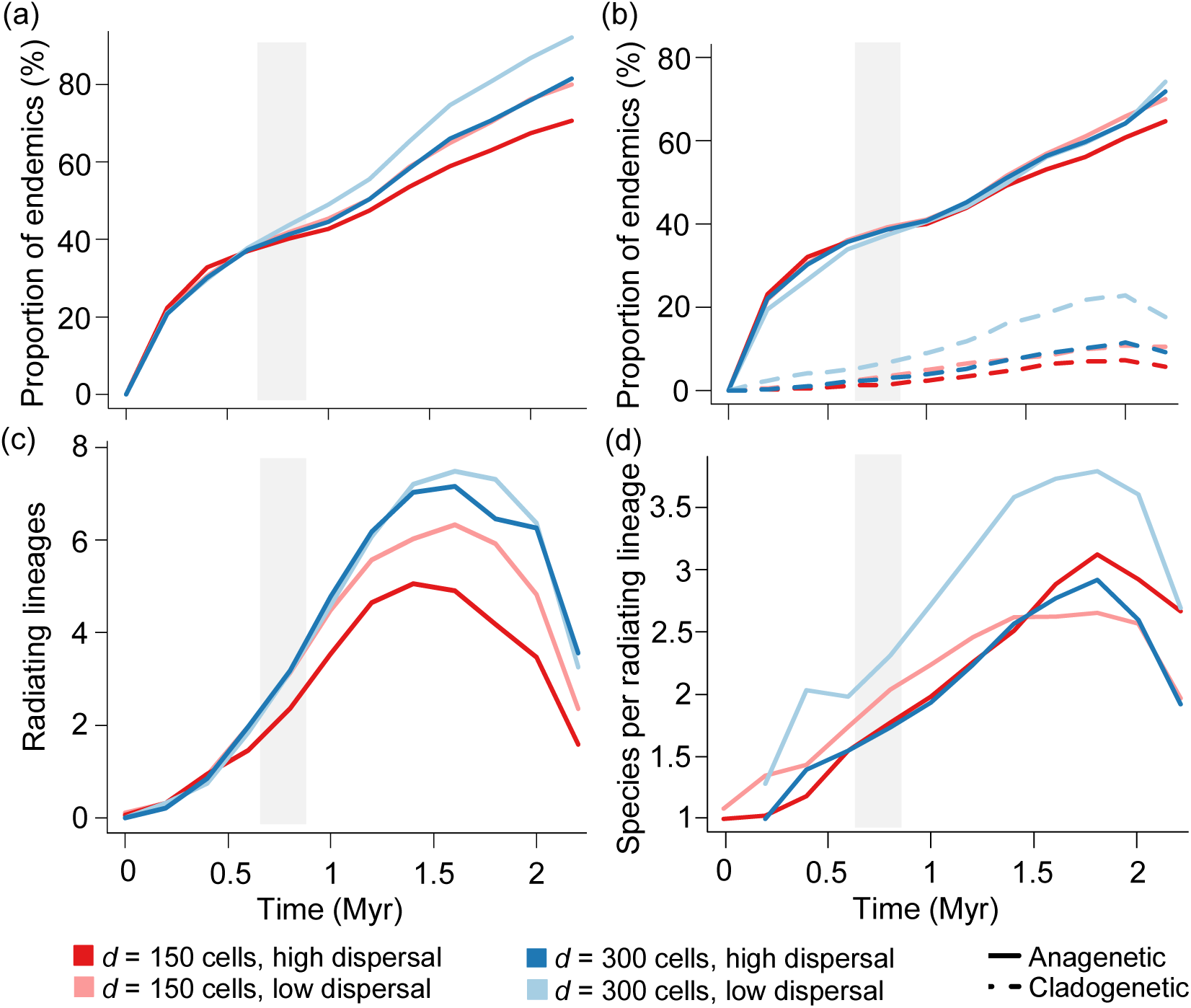
Speciation-related trends. (a) Proportion of all endemics. (b) Proportion of anagenetic and cladogenetic endemics (in each case as a function of all species). (c) Number of radiating lineages. (d) Number of species per radiating lineage. Note the steady increase in proportion of endemics in (a), mostly due to anagenetic endemics (b), despite overall humped richness trends shown in Fig. 2. Isolation scenarios were given by changing the distance *d* from mainland and long-distance dispersal ability of the source pool. Time-series were averaged within environmental time steps and over 20 replicate runs. The shaded area indicates the period with maximum island size.

Temporal trends in colonization rates were humped for all isolation scenarios, but the maximum values strongly decreased with isolation (Fig. 4a). Similar temporal trends but overall lower values were obtained for extinction rates (Fig. 4b). Anagenesis rates peaked at intermediate island age and monotonically decreased thereafter, with increasing isolation decreasing the maximum value (Fig. 4c). Cladogenesis rates were also humped and peaked at intermediate island age, with the amplitude of the curve increasing with isolation (Fig. 4d). Extinction rates of endemics increased over time and dropped sharply in the last thousand years (Fig. 4e). All islands showed a general positive net change rate (colonization rate + speciation rate – extinction rate) throughout the simulation (Fig. 4f), despite decreases in species richness during short-term peaks in extinction right after erosion events (averaged out across time steps in Fig. 4, but visible at higher temporal resolution as shown in Figure S2 of the Appendix S2). In our simulations, this increase was led by colonization rates, which were orders of magnitude higher than speciation rates.

**Figure 4.**
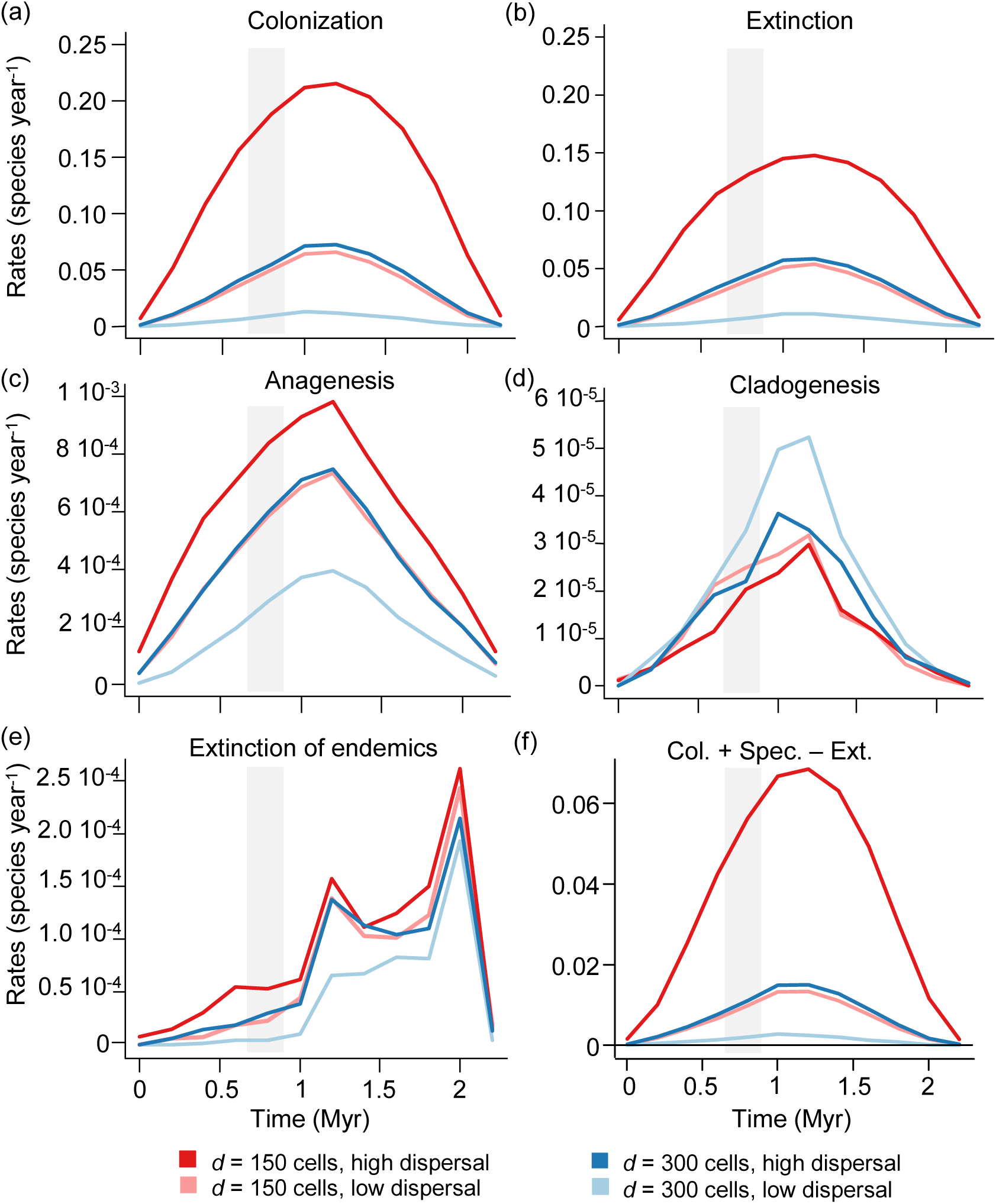
Biogeographical rates. (a) Colonization rates. (b) Extinction rates. (c) Anagenetic speciation rates. (d) Cladogenetic speciation rates. (e) Extinction rate of endemics. (f) Net rate: colonization + speciation - extinction rates. Isolation scenarios were simulated by changing the distance *d* from mainland and long-distance dispersal ability of the source pool. Rates are given in species per year, averaged within environmental time steps and over 20 replicate runs. The shaded area indicates the period with maximum island size.

## DISCUSSION

### Species richness

The simulation results for species richness patterns supported the GDM-derived hypotheses concerning temporal trends and isolation effects (Table 1). All isolation scenarios showed similar humped temporal trends in species richness, as predicted by the GDM based on the environmental dynamics of hotspot oceanic islands (Fig. 2a; Whittaker *et al.*, 2008; Borregaard *et al.*, 2016b; Cabral *et al.*, submitted) and from empirically well-supported, positive relationships between species richness and island area, elevation, and habitat heterogeneity (e.g. Kreft *et al.*, 2008; Hortal *et al.*, 2010). These three environmental factors positively affect species carrying capacity, a key concept in the GDM (Whittaker *et al.*, 2008). In our results, species richness seemed to approach carrying capacity only after area started to decrease. This was indicated by higher species richness during the erosion phase compared to the growth phase (Fig. 2a). This result is in line with the spatially-implicit model of Borregaard *et al.*, (2016b) but contrasts with indications for an early peak during island ontogeny (e.g. Steinbauer *et al.*, 2013; Lenzner *et al.*, 2016). Simulation experiments and empirical correlations must be compared with caution, as empirical estimations of carrying capacity entail shortcomings arising from the space-for-time substitution. For example, if the growth phase is several times shorter than the erosion phase, species richness is expected to peak relatively early. This should be particularly true if the island is within an archipelago, as immigration may be comparatively rapid in the youth of the island, due to colonization from the local (constrained) source pool provided by nearby islands. Here, BioGEEM simulates single islands and carrying capacity is an island property emerging from the environmental dynamics (in contrast of being a model parameter in Borregaard *et al.*, 2016b). These model features assure that how the carrying capacity changes and is filled does not depend on archipelagic effects but directly on available resources, environments, species properties, and on eco-evolutionary processes. This an advantage over spatially-implicit, neutral approaches, which often constrain carrying capacity via a parameter for the maximum number of individuals or species without considering individuals’ traits (Cabral *et al.*, 2016).

The fact that isolation consistently led to lower species richness over the entire island’s life span confirmed the GDM-based hypothesis (Table 1, Fig. 2a). Interestingly, both mechanisms of isolation (by distance and by dispersal ability) seemed interchangeable, as the scenarios with either isolation mechanism revealed very similar, intermediate values. Certain island models have also confirmed the negative effects of isolation on species richness, despite accounting for isolation only indirectly by varying colonization rates or island occupation at model initialization without explicitly considering geographic distances or species traits (Chen & He 2009; Rosindell & Phillimore, 2011; Rosindell & Harmon, 2013; Borregaard *et al.*, 2016b). These simpler approaches can simulate isolation effects, but provide limited interpretation of underlying ecological processes. In exploratory scenarios with alternative isolation mechanisms, BioGEEM also produced lower species richness with fewer mainland cells, dispersing species per time step, and species in the source pool (Figure S3 of the Appendix S2). Further modifications that remain to be tested involve processes at the intra-archipelagic level, including island hopping, parallel radiations, taxon cycles, and island merging (Cabral *et al.*, 2014; Weigelt *et al.*, 2016). This complexity in isolation and other eco-evolutionary mechanisms indicates that caution must be taken when interpreting results from correlations, as islands might be under the influence of different isolation mechanisms. It also offers a promising research avenue as most simulation models and correlative studies assume simplified isolation mechanisms (but see Weigelt & Kreft, 2013).

### Endemic richness

Our results for endemic richness supported the hypotheses on temporal trends for both anagenetic and cladogenetic endemics as well as isolation effects for cladogenetic endemics (Table 1). Endemic richness was humped and had a delayed peak compared to species richness (Fig. 2b; Steinbauer *et al.*, 2013; Borregaard *et al.*, 2016b; Cabral *et al.*, submitted). The opposing effects of isolation on anagenetic and cladogenetic richness (negative and positive, respectively) support previous findings of neutral models where considering islands that are already isolated (Rosindell & Phillimore, 2011). For weakly isolated islands, these neutral models demonstrated that intense gene flow from the mainland prevents both speciation modes (Rosindell & Phillimore, 2011). As isolation increases, reduced gene flow facilitates anagenesis. On already isolated islands, the further increasing isolation leads to a lower number of colonizing species, and thus a lower number of potentially anagenetic species. Simultaneously, radiating species start to fill empty niche space (Rosindell & Phillimore, 2011). This switch in predominant speciation mode emerges in neutral models simply by lowering colonization rates (Rosindell & Phillimore, 2011). While also capturing this switch, niche-based models explicitly integrate adaptive radiation and trait evolution (Cabral *et al.*, submitted). This is important for testing GDM predictions, as the GDM assumes that adaptive radiations occupy 'empty niches' (Whittaker *et al.*, 2008). In this sense, Cabral *et al.* (submitted) showed that radiating species tend to be ecologically distinct from cooccurring species, better occupying the niche space, as assumed by the GDM.

### Proportion of endemic species

Proportion of endemic species varied depending on speciation mode (Fig. 3), supporting the hypothesized isolation effects for all endemics and cladogenetic endemics as well as the temporal humped-trend only for cladogenetic endemics (Table 1). In our simulations, the overall endemism was mostly driven by anagenesis, which consistently increased over time but did not vary between isolation scenarios (Fig. 3b). The fact that the proportion of anagenetic endemics does not decrease indicates that anagenetic endemics are more likely to survive at the final island stage than cladogenetic endemics. This could arise because many anagenetic endemics evolved from colonizer species that were able to colonize and survive in such small islands and that did not establish radiating lineages. Thus, these anagenetic endemics have fewer close-competitors than species from cladogenetic lineages. Interestingly, Whittaker et al. (2008) also predict that the relative proportion of single-endemic lineages compared to multi-endemic lineages should increase over the final stages of an island’s existence, with one additional reason being the collapse of some multi-endemic lineages, ultimately to single representatives as the other members of former radiations become extinct.

In the final stages of island ontogeny, it is nevertheless expected within the GDM that proportion of single-island endemics drops (Whittaker *et al.*, 2008). This was based on two considerations. First it was argued that over time, some of these species colonize younger islands in an archipelago. Second, as islands become very old and low-lying islands (e.g. atolls), they may be increasingly characterized by disturbed, open, and strand-line habitats. These habitats facilitate the dominance of a few non-endemic, widespread, and disturbance-tolerant species, which make old islands with these habitats more akin to biologically “young” islands (see Dickinson 2009). BioGEEM does not simulate such conditions and it is thus unsurprising that anagenetic endemism did not decrease in our simulations. Nevertheless, the proportion of cladogenetic endemism showed the humped trend predicted by the GDM, but with a very delayed peak (Fig. 3b; Whittaker *et al.*, 2008). In real-world systems, the proportion of single-island endemics peaks at the same time as endemic richness (Steinbauer *et al.*, 2013). However, dispersal of single-island endemics to younger neighboring islands may contribute to earlier peaks, as some species lose their status as single-island endemics (see Borregaard *et al.*, 2016b).

Isolation effects varied the two different speciation modes, with no evident effect on the proportion of anagenetic endemics (also found by Stuessy *et al.*, 2006), but increasing the proportion of cladogenetic endemics (Fig. 3b). Hence, the observed higher proportion of all endemics on more isolated islands was largely driven by cladogenesis (Fig. 3a-b), confirming the important role of cladogenesis in filling empty niches on isolated islands, as assumed by the GDM premises (Whittaker *et al.*, 2008). In these premises, the higher prevalence of cladogenesis on remote, high-elevation islands seems more likely due to limited gene flow and greater ecological opportunities (Heaney, 2000; Stuessy *et al.*, 2006; Whittaker & Fernández-Palacios, 2007; Price & Wagner, 2011). Moreover, in their neutral and environmentally-static simulation analysis, Rosindell and Phillimore (2011) indicated a gradual replacement of anagenetic by cladogenetic endemics with increasing isolation. However, we did not obtain any replacement despite an increase in cladogenetic endemics (Fig. 3b). Instead, within the simulations, cladogenetic endemics seem to prevent colonization of non-endemics by being better adapted to local environments and communities than these naturally recurrent colonizers (Cabral *et al.*, submitted). Furthermore, only a minority of colonizing plant lineages on isolated islands are prone to diversification, while the majority of lineages only produce anagenetic endemics (compare Figs. 2b and 3c; Stuessy *et al.*, 2006; Price & Wagner, 2011). Such complex relationships between speciation modes can thus be addressed by integrating ecological, evolutionary, and environmental processes and by acknowledging species differences.

### Number of radiating lineages

Results for the number of radiating lineages supported the hypotheses on temporal trends and isolation effects (Table 1). Humped temporal trends were found in all scenarios as predicted by the GDM (Fig. 3c, Whittaker *et al.*, 2008), whereas isolation showed positive effects (Fig. 3c). Such results indicate that some lineages only radiate under low competition/colonization conditions. In fact, a few genera and families have been shown to be particularly radiation-prone wherever arriving in remote archipelagos, such as Hawaiian, Society, and Marquesa Islands (Price & Wagner, 2004; Price & Wagner, 2011; Lenzner *et al.*, 2016). Therefore, while knowledge about speciation on islands is constantly improving (e.g. Igea *et al.*, 2015), future simulation studies could investigate to what extent radiation-proneness is lineage-specific (e.g. mediated by common traits).

### Number of species per radiating lineage

Results for the number of species per radiating lineage were complex, supporting the hypothesis of an overall humped temporal trend (Fig. 3d; Whittaker *et al.*, 2008), but only partially supporting isolation effects, with an unexpected interaction between temporal and isolation effects (Table 1). Here, the isolation mechanism influenced the amplitude of the temporal trends, with an increase in species radiations becoming evident only for the most isolated islands (Fig. 3d). That is because islands isolated by just one isolation mechanism may still receive enough colonizers that occupy available niches. In contrast, islands isolated by both distance and dispersal can foster larger radiations than other islands due to less competition and greater ecological opportunities (e.g. empty niches; Whittaker & Fernández-Palacios, 2007). Therefore, large radiations may be more common where multiple isolation mechanisms are combined.

### Colonization rate

Results for the colonization rates supported the predicted negative isolation effects and predictions of humped temporal trends in all four isolation scenarios (Fig. 4a, Table 1). A decrease in colonization rates with isolation agrees with the predictions of both the equilibrium theory of island biogeography (MacArthur & Wilson, 1967) and the updated GDM (Borregaard *et al.*, 2016b). A higher rate of colonization on growing islands is consistent with an alleviated environmental filtering due to higher habitat heterogeneity and with a higher chance of a dispersal unit hitting the island (i.e. the target area effect – Lomolino, 1990). Environmental filtering and the target area effect are not explicitly considered by the GDM (Whittaker *et al.*, 2008). However, a simplified environmental filtering was considered by Borregaard *et al.* (2016b) by correlating colonization rates with carrying capacity.

### Extinction rate

Results for the extinction rates followed those of colonization rates, supporting hypothesized temporal and isolation effects (Fig. 4b, Table 1). When assessing only the extinction rates of endemic species, the hypotheses were also supported, but with a later peak and less difference between isolation scenarios (Fig. 4e, Table 1). The distinction between types of extinction is important. While endemic species can go globally extinct, non-endemics can (potentially) have their local extinction on an island reversed by re-colonization. In this sense, if colonization rates are much higher than speciation rates, extinction rates represent the extinction of non-endemics and reflect mostly colonization rate trends (Fig. 4a-b). If speciation rates have values comparable to or higher than colonization rates, such as in Borregaard *et al.* (2016b), extinction rates should reflect speciation rate trends. Furthermore, the overall positive net change rates (Fig. 4f; but see Figure S2 of the Appendix S2 for short, extinction-dominated net rate periods) indicate that a dynamic equilibrium cannot be achieved by the extinction rate that emerges in our experiments, namely from demographic stochasticity and environmental dynamics. In accordance, islands might steadily accumulate species if environmental dynamics are excluded (Cabral *et al.*, submitted). Future model developments might explore how extinction rates and the dynamic equilibrium vary with processes not implemented here, such as disturbances.

### Speciation rate

Results for speciation rates supported the hypotheses of temporal trends for both anagenesis and cladogenesis, but supported isolation effects only for cladogenetic endemics (Fig. 4c-d, Table 1). The differential effect of isolation on anagenesis and cladogenesis has not been the focus of the GDM, which focused predictions on species radiations, and thus cladogenesis (Whittaker *et al.*, 2008; Borregaard *et al.*, 2016b). In this sense, the increase in cladogenesis rates with isolation (Fig. 4d) was in accordance with empirical and modelling evidence (Heaney, 2000; Whittaker *et al.*, 2008; Rosindell & Phillimore, 2011; Borregaard *et al.*, 2016b). For the anagenesis rate, the negative isolation effect has been previously obtained by neutral models for islands that are already isolated enough to foster radiations (Rosindell & Phillimore, 2011). In BioGEEM, these differences between anagenesis and cladogenesis emerged within a niche-based approach, which indicates that disentangling the relative importance of neutral and non-neutral dynamics in real islands might not be trivial.

### Limitations and perspectives

The main limitation of BioGEEM is its complexity and data requirements for validation or parameterization (cf. Dormann *et al.*, 2012). Nevertheless, Cabral *et al.* (submitted) demonstrated that the model matches empirical and theoretical evidence at multiple ecological levels and that all simulated processes are necessary to simultaneously generate realistic patterns (i.e. 'pattern-oriented' modelling, *sensu* Grimm & Railsback, 2012). Therefore, scenario-based simulation experiments, such as presented here, have the potential to increase our understanding of process interactions and to generate simulation-driven hypotheses to be tested when appropriate data become available. Here, important differences compared to previous, simpler models are the explicit simulation of population-based processes (e.g. resource competition, stage transitions, individual dispersal) and species differences. These model features align with current trend in improving structural realism in ecological modelling (Cabral *et al.*, 2016; Grimm & Berger, 2016) and assure that biogeographical patterns are emergent system properties and not directly simulated via processes with biogeographical parameters (i.e. colonization, extinction, speciation rates as model parameters, not as emergent variables).

Another limitation is associated with the study design, namely that isolation did not vary over time. Isolation dynamics are not considered in the environmental dynamics assumed by the GDM and thus not accounted for in the present study. However, isolation changes at various time scales, from recently increased source pools via human-induced activities and dispersal (i.e. alien species; Kueffer *et al.*, 2010), over climate-mediated changes in sea levels over glaciation cycles (Fernández-Palacios *et al.*, 2015; Weigelt *et al.*, 2016), to deep-time scales of plate tectonics, archipelagic dynamics, and evolutionary changes in the source pool. Moreover, beyond the isolation mechanisms simulated in this study, intra-archipelagic isolation may play an important role for trends of single-island endemics (Cabral *et al.*, 2014; Borregaard *et al.*, 2016b), and, thus, further insights might be gained by disentangling intra-archipelagic isolation and connectivity and distance to different source pools. Integrative process-based frameworks, such as ours, provide excellent means to assess isolation dynamics explicitly, opening ground for future studies on theoretical developments and conservation assessments (e.g. due to human-induced sea level changes and alien species).

### Conclusions

In this simulation experiment, emergent patterns largely confirmed theoretical predictions of temporal trends and isolation effects on biogeographical patterns. However, noteworthy divergences that emerged included the steady increase in proportion of all endemics, as well as new insights regarding the extinction trends of endemics and the differential trends in proportion of endemics depending on speciation mode. Mechanistic simulation models like BioGEEM can thus contribute greatly to the theoretical understanding of the dynamics of complex systems. In fact, we demonstrated that dynamics of richness, endemism, colonization, speciation, and extinction emerge from individual- and population-level processes interacting with different isolation mechanisms. In the real-world, the dynamics of the insular environments, isolation, and area at the evolutionary time scale means that those ecological processes may never – or rarely – reach an equilibrium, which is reflected in our findings. Therefore, the adequate representation of persistent non-equilibrium conditions and the relevant processes affecting individuals and populations seems crucial to improving our understanding of biodiversity dynamics.

## ACKNOWLEDGEMENTS

J.S.C. acknowledges financial support from the German Research Foundation (DFG; SA 2133/11) and from sDiv, the Synthesis Centre of iDiv (DFG FZT 118). H.K. was supported by the DFG through the German Excellence Initiative. K.W. was partly funded by the State of Lower Saxony (Ministry of Science and Culture; Cluster of Excellence “Functional Biodiversity Research”). We thank Albert Phillimore, James Rosindell, Kostas Triantis, Yael Kisel, Gunnar Petter, Anke Stein, Patrick Weigelt, and Carsten Meyer for valuable feedback on a draft version of the manuscript.

## SUPPORTING INFORMATION

Additional supporting information may be found in the online version of this article:

**Appendix S1** Detailed model description.

**Appendix S2** Supporting figures.

## BIOSKETCH

**Juliano Sarmento Cabral** is interested in processes and factors influencing species and biodiversity dynamics across spatio-temporal scales. His research includes processes determining spatial and temporal distribution of tropical epiphytes, species ranges, island plant diversity as well as global species richness and endemism patterns.

## Author contributions

J.S.C. and H.K. designed the study, with input from K.W.; J.S.C. implemented and simulated the model; J.S.C. led the analyses and writing, with input from all coauthors.

